# Discovering Matrix Adducts for Enhanced Metabolite Profiling with Stable Isotope-Labeled MALDI-MSI

**DOI:** 10.1101/2023.06.28.546946

**Authors:** Gerard Baquer, Miguel Bernús, Lluc Sementé, René van Zeijl, Maria García-Altares, Bram Heijs, Omar Boutureira, Xavier Correig, Pere Ràfols

## Abstract

Matrix-assisted laser desorption ionization mass spectrometry imaging (MALDI-MSI) is a widely used technique for spatial metabolomics analysis, but the matrix introduces spectral interferences that impede data processing. In this study, we present an experimental and computational workflow utilizing isotopic labeling to discover and annotate matrix adducts in MALDI-MSI. Our approach enables the removal of matrix-related signals, improving metabolite annotation accuracy, extending metabolome coverage, and facilitating the interpretation of tissue morphology.

## 1. Introduction

Matrix-assisted laser desorption ionization mass spectrometry imaging (MALDI-MSI) has emerged as a powerful analytical technique for spatially resolving biomolecules in tissue samples^1^. While initially focused on large biomolecules like proteins and peptides^2, 3^, MALDI-MSI has recently gained attention in the study of small molecules such as lipids, metabolites, and drugs^4–6^. Proving invaluable in unraveling molecular networks in complex diseases such as cancer^7–10^.

However, the matrix used to facilitate desorption and ionization introduces spectral interferences in the low mass range, masking and suppressing target molecular species^11^, and posing challenges for untargeted metabolite profiling^12, 13^.

This problem can be mitigated with specialized sample preparation^14^ (e.g. derivatization, doping, or alternative deposition techniques) or with rationally designed MALDI matrices with minimal matrix interference (e.g. 1,5-diaminonaphthalene^15^, gold^16^, or silicon nanostructured substrates^17^). Nevertheless, first-generation matrices remain widely used due to their demonstrated analyte/matrix co-crystalization, spatial resolution, laser absorption, analyte ionization, and minimal analyte fragmentation. To date, 2,5-dihydroxybenzoic acid (DHB) represents 53% of all MALDI datasets in METASPACE^18^.

As an alternative, matrix signals can be annotated and removed computationally. rMSIcleanup^12^ uses spectral and spatial metrics to annotate matrix signals in Ag-LDI-MSI. Mass2adduct^19^ exploits mass differences between all possible pairs of ions to find metabolite-matrix adducts. And OffSampleAI^20^ uses machine learning (ML) to annotate ion images localized outside of the tissue. Current methods are limited as they (1) rely on a predefined list of matrix adducts, (2) assume a matrix-like spatial distribution, (3) fail to control the false discovery rate (FDR), and (4) lack comprehensive validation and characterization of organic matrix adduct formation.

To address this challenge and enhance metabolite profiling, we present an experimental and computational workflow that exploits stable isotope-labeled (SIL) MALDI-MSI to discover matrix adducts. Using a DHB analog with a ^13^C-labeled aromatic ring (^13^C_6_-DHB) synthesized in-house we effectively classify MS features into analytes, contaminant-matrix adducts, and analyte-matrix adducts. Matrix-containing MS features are then annotated (with target-decoy FDR estimation^21^) against a theoretical list of DHB adducts, in-source fragments, and metabolites (HMDB^22^).

We show that matrix signals represent 17.7% of all ions (SNR>5) and their removal is imperative in untargeted applications, as it improves metabolite annotation, dimensionality reduction, and spatial clustering. Additionally, 16% of all ions correspond to analyte-matrix adducts, so searching for analyte-matrix adducts increases the metabolome coverage of untargeted MALDI-MSI analyses.

## 2. Results

### 2.1. Synthesis of High-Purity ^13^C_6_-DHB

The synthesis of ^13^C_6_-DHB was designed using ^13^C_6_ labeled salicylic acid as starting material, given its commercial availability in high isotopic enrichment, and its structural similarity to the target molecule. The synthetic sequence involved two steps (Figure 1): (1) selective electrophilic bromination of the 5^th^ position with N-bromosuccinimide in acid media and, (2) Cu-mediated Ullmann-type hydroxylation. This sequence had a moderate yield of 49% which was later purified by flash SiO_2_ chromatography to obtain high-purity ^13^C_6_-DHB (>99%, determined by HPLC).

**Figure 1.**
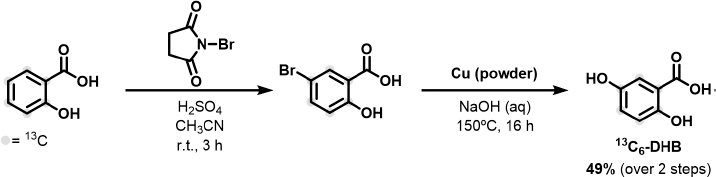
Synthesis of ^13^C_6_-DHB by selective electrophilic bromination of the 5^th^ position with N-bromosuccinimide in acid media followed by Cu-mediated Ullmann-type hydroxylation.

### 2.2. ^13^C_6_-DHB Produces High-Quality MALDI-MS Images

In this section, we compare the quality of the data produced with DHB (n=2) and ^13^C_6_-DHB (n=2) to ensure that ^13^C_6_-DHB can be used for MALDI-MSI (Figure 2). To block biological variability, we used 4 consecutive sagittal sections of a single mouse brain.

**Figure 2.**
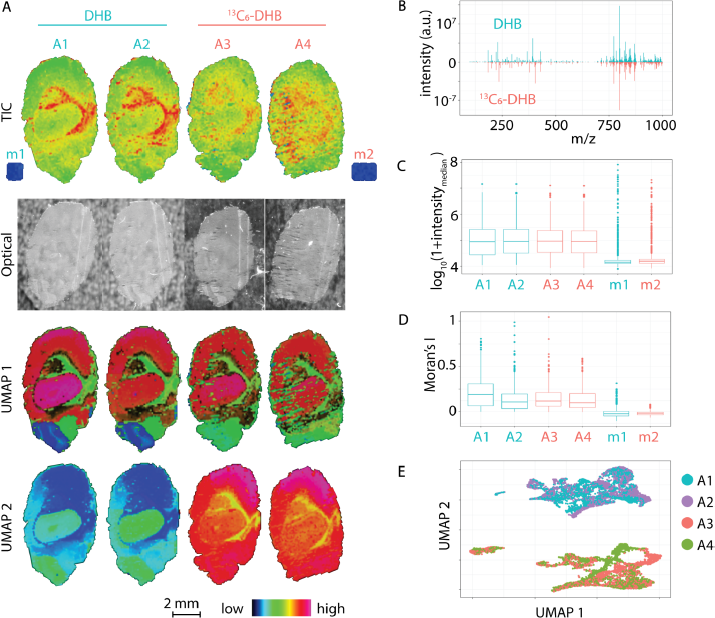
DHB vs ^13^C_6_-DHB MALDI-MSI data quality comparison. **(A)** Spatial visualization of single-spectrum total ion count (TIC) values. **(B)** Mean spectra of DHB vs ^13^C_6_-DHB samples. **(C)** Median intensity per MS feature. **(D)** Morańs I autocorrelation per MS feature as a measure of morphological definition. **(E)** Optical images of the DHB and ^13^C_6_-DHB sections. **(F)** UMAP projection of all recorded MSI pixels based on their individual spectra. **(G)** Spatial visualization of UMAP 1. **(H)** Spatial visualization of UMAP 2.

In all samples, spectra recorded from the corpus callosum present higher total ion count (TIC) (Figure 2A first row), commonly attributed to its dissimilar ion suppression compared to densely packed brain tissue^23^. The mean spectra have comparable intensities and MS features (Figure 2B) and the distribution of pixel TIC across DHB and ^13^C_6_-DHB samples is identical (Figure 2C).

To assess the spatial quality of the data, we rely on spatial autocorrelation^24^ (Moran’s I) as a measure of the morphological definition of each ion. Figure 2D shows that matrix control regions, specifically recorded off-sample regions (only matrix), show negligible autocorrelation given their spatial homogeneity and noise. In contrast, tissue regions present positive autocorrelation suggesting a fine definition of morphological features. Sample A1 (DHB) presents higher autocorrelated ions, indicating a better definition of anatomical structures. Nevertheless, this difference is not statistically significant (*p*-value = 0.83) when compared to ^13^C_6_-DHB samples. Optical images (Figure 2A second row) reveal more sectioning artifacts (cracks and folds) in the ^13^C_6_-DHB samples and explain the slightly lower-quality MS images.

Finally, a UMAP^25^ projection of all pixels based on their MS spectra (Figure 2E) reveals exceptionally low variability among technical replicates and clear separation between DHB and ^13^C_6_-DHB. When mapped spatially, the first component (UMAP 1) (Figure 2A third row) is similarly expressed regardless of the matrix used and highlights different anatomical features: cerebral cortex and midbrain (red), cerebral nuclei (black), thalamus and hypothalamus (pink), corpus callosum (green) and medulla (blue). On the other hand, the second component (UMAP 2) (Figure 2A fourth row) highlights differences between DHB and ^13^C_6_-DHB. These differences are expected; while the same matrix-related molecules are present in all samples their associated *m/z* signals are different due to the introduced *m/z* shift. Only analyte ions are shared.

In summary, our analyses collectively demonstrate that ^13^C_6_-DHB MALDI-MSI yields high-quality ion images, presenting a promising avenue for discovering matrix signals.

### 2.3. ^13^C_6_-DHB Enables the Discovery of Matrix Signals

#### 2.3.1. Workflow Overview

The workflow for matrix signal discovery is summarized in Figure 3. Four consecutive tissue sections are sprayed with DHB (n=2) and ^13^C_6_-DHB (n=2). To identify matrix-related signals, we search for MS feature pairs (DHB vs ^13^C_6_-DHB) with specific theoretical *m/z* shifts (e.g. +6, +12, and +18 Th) and matching spatial distributions. These are further split into analyte and contaminant adducts based on their presence in off-sample regions (only matrix). Our method effectively classifies all ions into (1) analytes, (2) contaminant-matrix adducts, and (3) analyte-matrix adducts (Figure 3A).

**Figure 3.**
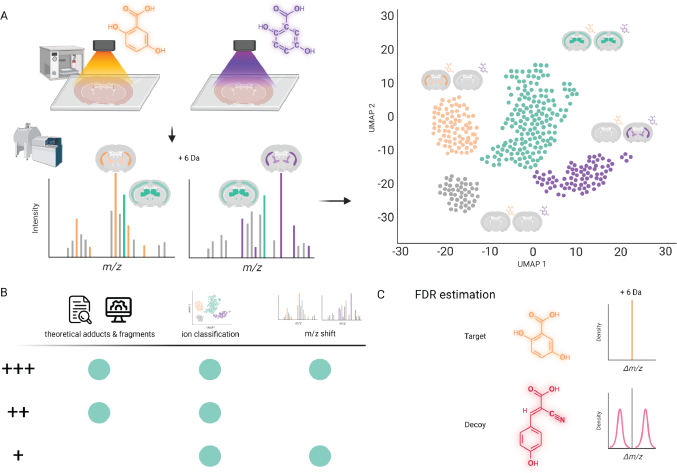
General workflow of matrix adduct discovery using SIL-MALDI-MSI. **(A)** Experimental and computational foundations. **(B)** Annotations are split into 3 confidence levels based on the evidence found. **(C)** FDR-estimation based on a decoy matrix and a decoy bimodal distribution of *m/z* shifts.

Analyte signals should be annotated with external tools (e.g. METASPACE^18^, LipostarMSI^26^, or rMSIannotation^27^) while matrix signals are annotated against a database containing theoretical DHB adducts, in-source fragments, and metabolites (HMDB^22^). The annotations are assigned a confidence level based on the evidence found (theoretical match, spatial classification, and SIL *m/z* shift) (Figure 3B). The top two levels (+++ and ++) correspond to hits against the database (see Methods and Supplementary Table 8) while the bottom level (+) represents candidate adducts of DHB with analytes not found in the database. Additionally, we estimate the false discovery rate (FDR) in a target-decoy setting^28^, using a decoy matrix (CHCA) and bimodal distribution of decoy *m/z* shifts (Figure 3C).

#### 2.3.2. Exploiting Labeling-Induced *m/z* Shifts

To gauge the potential of leveraging labeling-induced *m/z* shifts for matrix annotation, we explore the most abundant *m/z* shifts between the DHB and ^13^C_6_-DHB samples (Figure 4A). The three most prevalent shifts (z-score > 2**σ**) include analyte signals (+0 Th), M+1 isotopes (+1 Th), and [^13^C_6_-DHB - DHB] (+6 Th). Another notable shift (z-score > **σ**) is the 2·[^13^C_6_-DHB - DHB] *m/z* shift (+12 Th). Figure 4B demonstrates the absence of labeling-induced m/z shifts (+6 and +12 Th) comparing the two DHB replicates, serving as a negative control.

**Figure 4.**
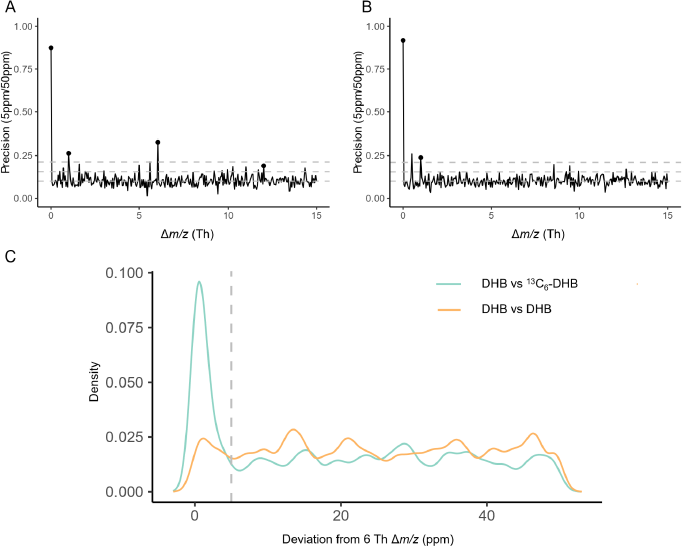
Exploration of *m/z* shifts **(A)** Precision of *m/z* shifts in DHB vs ^13^C_6_-DHB **(B)** Precision of *m/z* shifts in DHB vs DHB **(C)** Density distribution of deviation from +6 Th *m/z* shift (ppm).

To further validate the precision of labeling-induced shifts, we asses the distribution of shifts around +6 Th (ε<50ppm) (Figure 4C). A large portion of the shifts fall within a 5 ppm threshold, indicating high precision (AUC_5ppm_/AUC_50ppm_=0.34). In contrast, comparing the two DHB samples yields a uniform distribution with baseline precision (AUC_5ppm_/AUC_50ppm_=0.1). As a negative control, a non-labeling-induced *m/z* shift (+2 Th) exhibits unspecific uniform distributions in both comparisons (AUC_5ppm_/AUC_50ppm_=0.1) (Supplementary Figure 1).

To sum up, labeling-induced m/z shifts are a precise metric to screen for potential matrix signal candidates.

#### 2.3.3. Leveraging Spatial Similarity

To refine the list of potential matrix signal candidates, we analyzed their spatial distribution. First, we facilitate cross-sample spatial comparison by bringing all four samples into the same coordinate system using image registration (see Methods). We calculated the following metrics for each ion: (1) spatial correlation across all samples, (2) spatial correlation within DHB samples, (3) spatial correlation within ^13^C_6_-DHB samples, and (4) mean intensity in each matrix control. These metrics were then visualized on a UMAP projection, which represented the spatial similarity between all MS features (Figure 5A). The three spatial correlations designate distinct regions in the projection corresponding to analytes, DHB, and ^13^C_6_-DHB (Figures 5B, C & D). By considering the mean ion intensity in the matrix control regions, we further differentiated each matrix-related group into on-sample and off-sample (Figures 5E & F). Ions that did not meet the specified criteria were considered background, as they provided minimal spatial information and had a signal-to-noise ratio (SNR) below 5.

**Figure 5.**
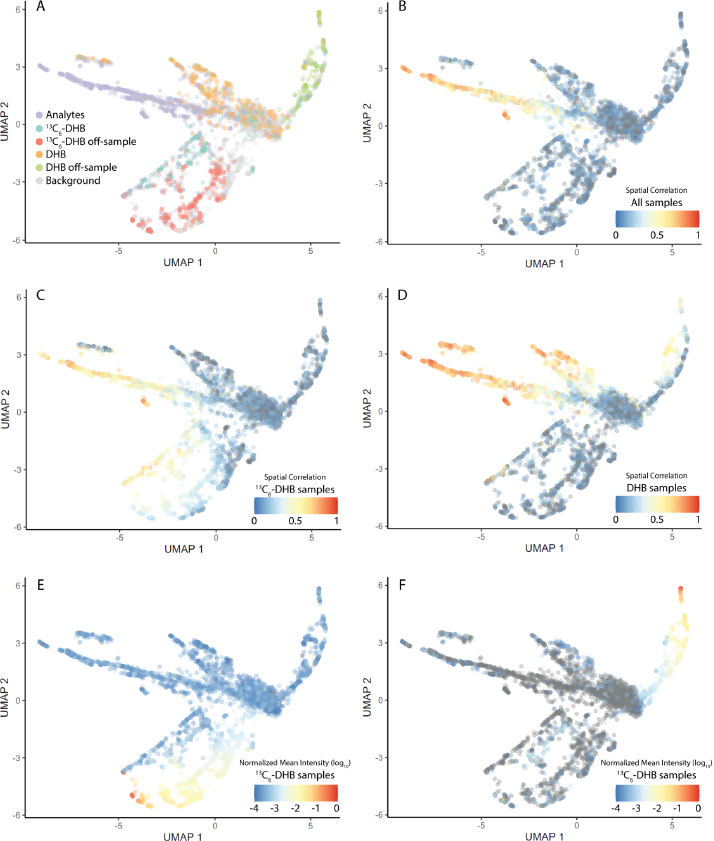
Classification of ion images based on their spatial distribution. **(A)** Classification of each ion image. **(B)** Spatial correlation across all samples, **(C)** ^13^C_6_-DHB samples, and **(D)** DHB samples. **(E)** Mean ion intensity in DHB off-sample region and **(F)** ^13^C_6_-DHB off-sample region.

Figure 6 provides a visual representation of spatial correlations among representative ions from the three main classes. Ions *m/z* 203.2233 and *m/z* 190.0122 exhibit strong spatial correlation across all samples. On the other hand, DHB on-sample ions like *m/z* 213.9642 show a high spatial correlation only within DHB samples, while ^13^C_6_-DHB on-sample ions like *m/z* 219.9839 only show a high spatial correlation within ^13^C_6_-DHB samples. Notably, these two ions correspond to the same matrix-related signal as indicated by the +6 Th *m/z* shift and their high spatial correlation (r=0.31, *p*-value < 0.01).

**Figure 6.**
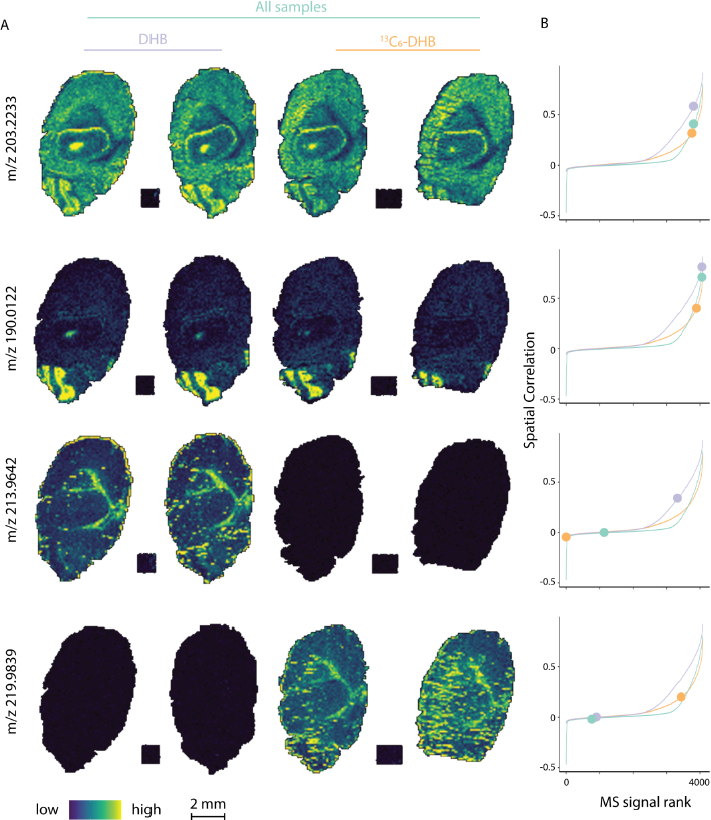
Representative examples of ion classification based on their spatial distribution. **(A)** Ion images and **(B)** rank order plot of spatial correlation and ion images for *m/z* 203.2233, *m/z* 190.0122, *m/z* 213.9642, and *m/z* 219.9839

In conclusion, image registration and cross-sample correlation improve matrix signal annotation and enable their accurate classification into on-sample and off-sample.

#### 2.3.4. Discovered Matrix Adducts

Using the described protocol, we annotated twelve matrix-contaminant adducts (Table 1) and uncovered 112 m/z unidentified features that formed adducts with the matrix (Supplementary Table 6). By cross-referencing with the HMDB, we obtained 59 metabolite-matrix annotations for these m/z features (Supplementary Table 7).

**Table 1.**
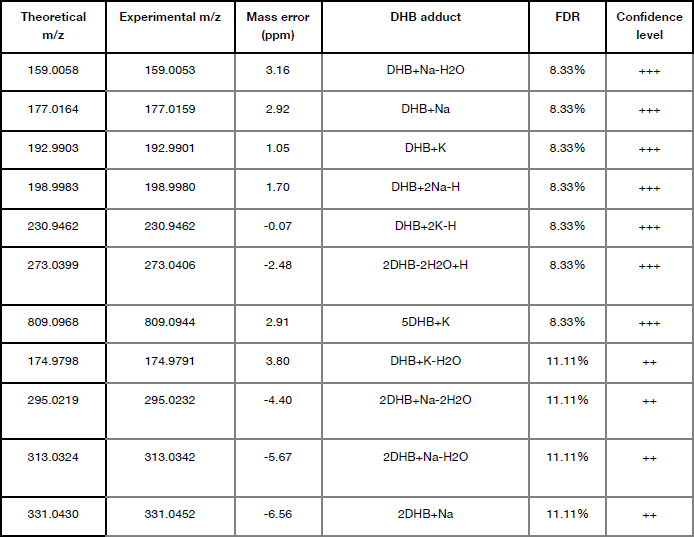
List of discovered contaminant DHB adducts. The False Discovery Rate (FDR) is estimated using a target-decoy approach (see Methods & Figure 3C). The confidence levels are based on the available evidence (see Methods Figure 3B).

| Theoretical m/z | Experimental m/z | Mass error (ppm) | DHB adduct | FDR | Confidence level |
| --- | --- | --- | --- | --- | --- |
| 159.0058 | 159.0053 | 3.16 | DHB+Na-H<sub>2</sub>O | 8.33% | +++ |
| 177.0164 | 177.0159 | 2.92 | DHB+Na | 8.33% | +++ |
| 192.9903 | 192.9901 | 1.05 | DHB+K | 8.33% | +++ |
| 198.9983 | 198.9980 | 1.70 | DHB+2Na-H | 8.33% | +++ |
| 230.9462 | 230.9462 | -0.07 | DHB+2K-H | 8.33% | +++ |
| 273.0399 | 273.0406 | -2.48 | 2DHB-2H<sub>2</sub>O+H | 8.33% | +++ |
| 809.0968 | 809.0944 | 2.91 | 5DHB+K | 8.33% | +++ |
| 174.9798 | 174.9791 | 3.80 | DHB+K-H<sub>2</sub>O | 11.11% | ++ |
| 295.0219 | 295.0232 | -4.40 | 2DHB+Na-2H<sub>2</sub>O | 11.11% | ++ |
| 313.0324 | 313.0342 | -5.67 | 2DHB+Na-H<sub>2</sub>O | 11.11% | ++ |
| 331.0430 | 331.0452 | -6.56 | 2DHB+Na | 11.11% | ++ |

### 2.4. Generalizability across Multiple DHB Datasets

To validate the DHB adducts, we annotated consecutive mouse brain sections and off-sample matrix controls prepared with DHB, Au, 9AA, NEDC, and Norharmane (Figure 7 & Supplementary Tables 1, 3 & 4). Most DHB adducts (confidence +++ or ++) were detected in all DHB samples, but potassium adducts were absent. Adducts [2DHB+Na-2H_2_O]^+^ and [2DHB+Na]^+^ were exclusively detected in off-sample regions, while [DHB+2K-H]^+^ was only found on tissue. The absence of DHB adducts in non-DHB samples demonstrates high specificity (Figure 7A).

**Figure 7.**
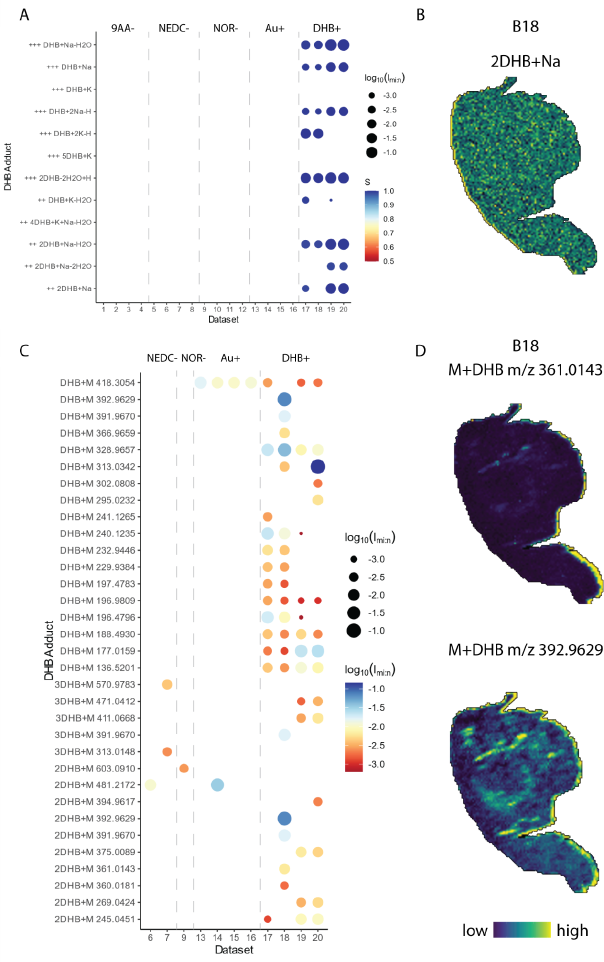
Annotation of mouse brain samples prepared with different matrices. (A) Contaminant-matrix-related annotations (+++ and ++). The color indicates the S score (see Methods) and the size indicates the intensity normalized to the maximum peak in the mean spectrum. **(B)** Example ion images. **(C)** Analyte matrix-related annotations (+), and **(D)** example images.

While lower-confidence annotations with unassigned formulas (+) were comprehensively covered in DHB samples, there were occurrences in non-DHB datasets, suggesting lower specificity (Fig. 7C). As notable DHB-specific examples: m/z 197.4783 and 232.9446 were exclusively detected on-sample, m/z 269.0424, 375.0089, and 471.0412 were exclusively found off-sample, while m/z 136.5201 and 188.4930 were present in both cases. The main unspecific peak, *m/z* 418.3054, is consistently detected in all Au samples, suggesting it could be a metabolite ionized by both matrices.

We conducted the same annotation on human biopsies of diverse tissues (brain, lung, kidney, and liver) prepared with DHB (Supplementary Table 2). All samples expressed several DHB-related adducts with some tissue-specific differences (Figure 8A). The coverage in lung and kidney samples (#1-#6) was considerably lower than in the brain samples (#9-#13). This could be explained by the higher sodium levels in the brain^29^.

**Figure 8.**
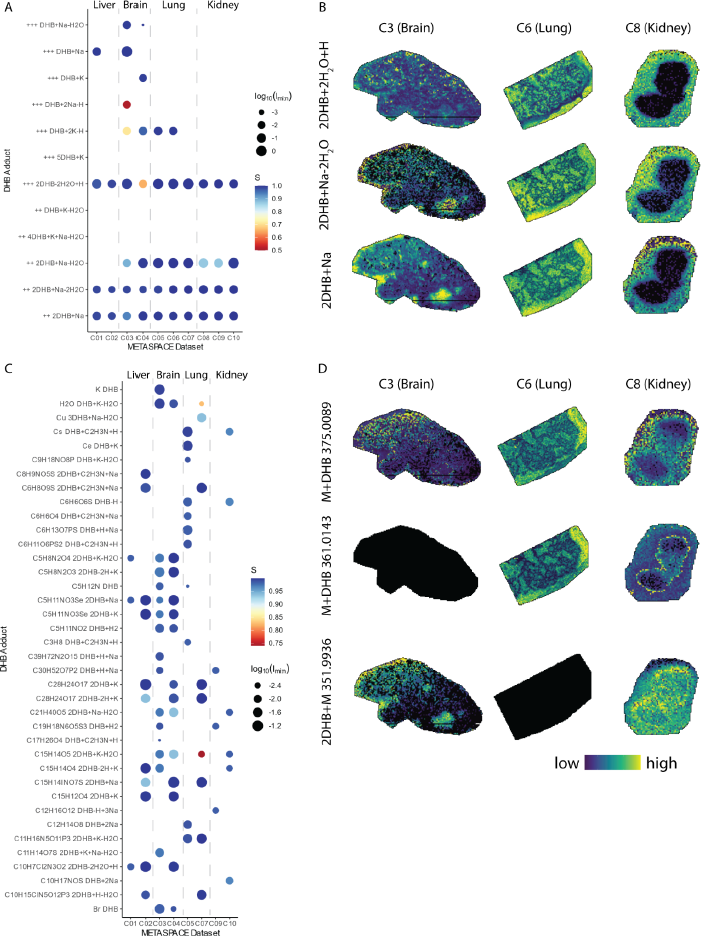
Annotation of human biopsies of different organs prepared with DHB (METASPACE). **(A)** Contaminant-matrix-related annotations (+++ and ++). The color indicates the S score and the size indicates the intensity normalized to the maximum peak in the mean spectrum. **(B)** Example ion images. Gray images indicate the given ion is not present in the sample. **(C)** Analyte-matrix-related annotations matched against HMDB (+++). **(D)** Example ion images of analyte-matrix-related annotations that did not return any hits against HMDB (+).

Figure 8C shows the annotation of DHB adducts with analyte molecules matched against HMDB. As an example, we find the following adduct C_21_H_40_O_5_ + 2DHB + Na - H_2_O in all brain samples and one kidney sample. HMDB annotates this molecule as a diacylglycerol (DG) isomer with 0, 8, and 10-carbon saturated fatty acid chains. DGs are involved in various physiological processes, including cell membrane formation, energy storage, and intracellular signaling pathways^30^. They can be synthesized in the body or obtained from dietary sources, and have been found in the brain, where they are implicated in neural signaling and synaptic function^30^.

In both validations, ion images of contaminant-matrix adducts show homogeneous or off-tissue distributions (Figure 7B & 8B) while analyte-matrix adducts present biologically-relevant morphologies (Figure 7D & 8D).

In summary, the annotated matrix adducts are unique to DHB and can be detected in various clinical samples.

### 2.5. Removal of Matrix Signals Improves Post-Processing

#### 2.5.1. Effects of Matrix Removal on Dimensionality Reduction

In this section, we explore the influence of matrix-related peaks in a typical untargeted analysis including dimensionality reduction and small molecule annotation.

Firstly we conduct a UMAP dimensionality reduction of all pixels in all consecutive mouse slices prepared with DHB and ^13^C_6_-DHB. As discussed in the quality control section, UMAP mainly highlights the difference between the two matrices (Figure 9A) while the technical replicates within each group are closely intertwined. After the removal of all matrix-related peaks, the two groups are brought closer together and are almost indistinguishable (Figure 9B). These differences are better understood when contextualized in the spatial context (Figure 9C). When using all peaks in the samples both UMAP projections highlight differences between groups and convey few morphological features. After removing matrix-related signals both projections highlight different anatomical features with high contrast and are almost indistinguishable irrespective of DHB labeling. The only apparent difference is the definition of the cortex in UMAP 2. These findings indicate correct and comprehensive annotation of matrix-related signals.

**Figure 9.**
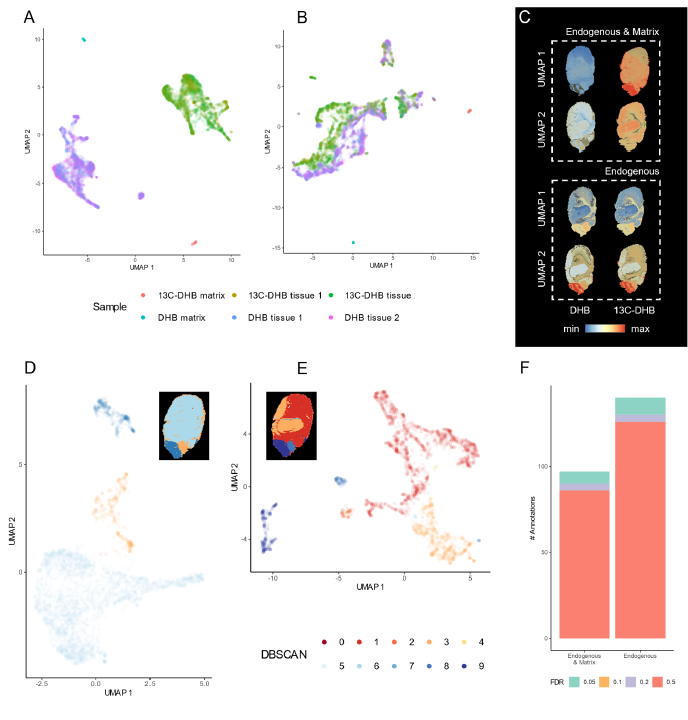
Influence of matrix-related signals in untargeted workflows DHB samples. **(A)** UMAP pixel projection before and **(B)** after matrix removal **(C)** Spatial UMAP mapping before (top) and after (bottom) matrix removal **(D)** UMAP pixel projection and DBSCAN clustering before, and **(E)** after matrix removal **(F)** Number of METASPACE annotations before and after matrix removal.

Similarly, when only analyzing Sample #1 (DHB), UMAP can better capture morphological features after the removal of matrix-related signals (Figures 9D & E). DBSCAN clustering ^31^ of the projection with matrix-related peaks is only capable of identifying 3 major clusters (Figure 9D) (background, cerebellum, and cerebrum+brain stem). When exclusively using analyte ions, the same clustering identifies at least 5 major regions (background, cerebellum, cortex+interbrain, nuclei+midbrain, and hindbrain), (Figure 9E).

#### 2.5.2. Effects of Matrix Removal on Metabolite Annotation

In the second part of our untargeted analysis, we perform metabolite annotation (METASPACE + HMDB)^18, 22^ on Sample #1, both before and after eliminating matrix-related signals (Figure 9F). With all signals retained, METASPACE returned 97 annotations, with only 7 highly-reliable annotations (FDR<5%). However, removing matrix-related signals resulted in an increased number of annotations, reaching 126, with 10 highly-reliable annotations. When comparing the annotated molecular formulas, the analyte-only dataset retains 86% of annotations and 100% of reliable annotations (FDR<20%) from the complete dataset. Our findings demonstrate that removing matrix-related signals reduces decoy hits, preserves target hits, lowers FDR values, and enhances the overall confidence of annotation.

## 3. Discussion

We propose an innovative workflow using SIL-MALDI-MSI to enable untargeted metabolomics. By synthesizing ^13^C_6_-DHB and exploiting their known m/z shift and preserved spatial distribution we can annotate all matrix signals. To ensure confident annotation we introduce a novel FDR estimation paradigm based on decoy matrices and *m/z* shifts.

Using this approach, we show that matrix signals represent 17.7% of all ions (SNR>5) of which 90% (16% of all ions) correspond to analyte-matrix adducts. The resulting theoretical list of matrix-related signals can be used as a library to annotate matrix signals in MALDI-MSI experiments with comparable conditions (i.e., tissue type, matrix, and ionization polarity). Different matrices and conditions would require a new pilot study using SIL-MALDI-MSI and the computational workflow presented to discover matrix signals.

This work reveals a crucial insight: matrix signals negatively affect MALDI-MSI untargeted metabolomics. By removing these signals, dimensionality reduction algorithms like UMAP^25^ can better emphasize biologically and anatomically relevant structures. Additionally, excluding matrix signals improves the annotation of small molecules using automated tools like METASPACE^18^. This removal results in a higher number of annotations with increased confidence (lower FDR). Furthermore, we demonstrate that 16% of all ions correspond to matrix adducts with analytes and their annotation expands metabolite coverage significantly.

In conclusion, SIL matrices resolve a major bottleneck MALDI-MSI metabolomics by comprehensively annotating matrix signals. This enhances untargeted analyses, boosts confidence in metabolite annotation, and expands metabolome coverage.

## Methods

### 2,5-dihydroxybenzoic-1,2,3,4,5,6-^13^C_6_ acid, (^13^C_6_-DHB) synthesis

In a 25 ml round bottom flask equipped with a magnetic stir bar, 2-hydroxybenzoic-1,2,3,4,5,6-^13^C_6_ acid (855.7 mg, 5.94 mmol) was dissolved in 12 ml of acetonitrile (0.5 M). Next, concentrated sulfuric acid (334 µl, 6.23 mmol, 1.05 eq.) was added dropwise and the solution was stirred for 5 minutes. Then, N-bromosuccinimide (1.11 g, 6.23 mmol, 1.05 eq.) was added and the mixture was left stirring at room temperature for 3 hours. Reaction completion was determined by TLC analysis. The mixture was evaporated under reduced pressure, redissolved with water, and extracted with CH_2_Cl_2_ (3 x 15 ml). The combined organic layers were washed brine, dried over MgSO_4_, and concentrated under reduced pressure to afford the crude product (1.428 g) which was used directly in the next step.

In a pressure Fisher-Porter reactor equipped with a magnetic stir bar, 300 mg of the crude mixture from the previous reaction was dissolved in 7.1 ml of an aqueous solution of NaOH (8%). Then, freshly activated copper powder (257 mg, 4.04 mmol, 3 eq.) was directly added to the vessel. Next, the reaction was heated up to 150°C and stirred for 16 h. After cooling the reactor to room temperature, the crude mixture was filtrated through celite and acidified with HCl (aq. 10%), and extracted with Et_2_O (3 x 15 ml). The combined organic layers were washed brine, dried over MgSO_4_, and concentrated under reduced pressure. The crude was then purified by flash chromatography to obtain the desired 2,5-dihydroxybenzoic-1,2,3,4,5,6-^13^C_6_ *acid* (105.8 mg) as a pale brown solid in 49% yield.

### ^13^C_6_-DHB purity tests

Proton (^1^H NMR) and carbon (^13^C NMR) nuclear magnetic resonance spectra were recorded on a Varian Mercury spectrometer (400 MHz for ^1^H) and (100.6 MHz for ^13^C). All chemical shifts are quoted on the *δ* scale in parts per million (ppm) using the residual solvent as the internal standard (^1^H NMR: CD_3_OD = 3.31 and ^13^C NMR: CD_3_OD = 49.0). Coupling constants (*J*) are reported in Hz with the following splitting abbreviations: s = singlet, d = doublet, t = triplet, q = quartet, and m = multiplet. Thin layer chromatography (TLC) was carried out using commercial-backed sheets coated with 60 Å F_254_ silica gel. Visualization of the silica plates was achieved using a UV lamp (λ_max_ = 254 nm) and potassium permanganate staining solutions. Flash column chromatography was carried out using silica gel 60 Å CC (230–400 mesh). Mobile phases are reported in relative composition (*e.g.* 1:1 Ethyl acetate/hexane v/v). Brine refers to a saturated solution of sodium chloride. Anhydrous magnesium sulfate (MgSO_4_) was used as a drying agent after the reaction work-up, as indicated. All reagents were purchased from Sigma Aldrich, Cymit, Carbosynth, Apollo Scientific, Fluorochem, and Eurisotop chemical companies.

**^1^H NMR (CD_3_OD, 400 MHz):** δ 7.49−7.01 (m, 1H), 7.20−6.71 (m, 1H), 7.01−6.52 (m, 1H).

**^13^C{^1^H} NMR (CD_3_OD, 100.6 MHz):** δ 157.2−155.7 (m), 152.6−148.0 (m), 124.7 (ddt, J = 65.0, 60.3, 4.9 Hz), 118.7 (ddt, J = 68.4, 60.3, 4.7 Hz), 117.2−114.7 (m), 114.6−112.3 (m).

**HRMS (MALDI-FTICR-MSI):** for (M+Na)^+^ C^13^C_6_H_6_NaO ^+^ (m/z): calculated 183.0360; found 183.0366.

**Purity (HPLC):** >99%.

### Sample Preparation and MALDI-MSI Acquisition

The sample preparation and acquisition parameters for all the samples used in this study are summarized in Supplementary Tables 1 and 2.

For samples A1-A4 and B1-B20 mice brains were collected from C57BL/6 mice, fresh-frozen in liquid nitrogen, and stored at −80°C. Brains were sectioned in a cryo-microtome (Leica Microsystems, Wetzlar, Germany) set to −20°C to obtain 10 µm-thick tissue sections, which were thaw-mounted on indium-tin-oxide (ITO)-coated glass slides (Bruker Daltonics, Bremen, Germany), and again stored at −80°C until further use. Prior to analysis, slides were equilibrated to room temperature in a vacuum freeze-drier for 20 minutes and coated with the MALDI matrix. Prior to matrix application, the matrix was checked for contaminants by spotting the matrix on a target plate and profiling the spots. The compositions for the different matrices can be found in Supplementary Table 3, and the SunCollect (SunChrom GmbH, Friedrichsdorf, Germany) application methods used for the different matrices can be found in Supplementary Table 4.

During the definition of the measurement areas both tissue sections, as well as an off-tissue matrix area, were defined for measurement. The off-tissue matrix areas were used to distinguish between on-tissue and off-tissue matrix-specific *m/z* features.

MALDI-FTICR-MSI was performed on two solariX (Bruker Daltonics) platforms, both equipped with a CombiSource. The system used for samples A1-A4 was equipped with a 12 T superconductive magnet. Samples B1-B20 were acquired using a 9.4 T superconductive magnet. The parameters used for the different measurements can be found in Table xx. In short, all datasets were recorded over an *m/z* range between 100-1000 Th, using 100×100 µm^2^ pixels and a 512k datapoint transient. The measurement polarity, laser intensity, number of shots per pixel, and laser frequency were selected for each matrix (Supplementary Table 5). All methods were externally calibrated using red phosphorous as a mass calibrant. Data were acquired using ftmsControl (v2.1; Bruker Daltonics) and flexImaging (v5.0; Bruker Daltonics).

### MSI data processing

All samples were visualized and exported to .imzML using SCiLS (Bruker). The .imzML files were processed and exported to a centroid-mode peak matrix using rMSIproc^32^. The default processing parameters were used. The Signal-to-Noise Ratio (SNR) threshold was set to 5 and the Savitzky–Golay smoothing had a kernel size of 7. Peaks appearing in less than 5% of the pixels were filtered out. Peaks within a window of 6 data points or scans were binned together as the same mass peak. No intensity normalization was performed.

Image registration of Datasets A1-A4 was performed using rMSIworkflows (https://github.com/gbaquer/rMSIworkflows). Manually specified fiducial markers were used to calculate and apply a rigid transformation (rotation, translation, and scaling). The pixel coordinates of all samples were aligned into the same coordinate space to enable spatial correlation of the same metabolite across samples.

### Statistical methods

All statistical group comparisons were performed at a pixel level using the linear mixed effects model in the “nlme” R package^33^. We considered sample ID as a random effect and adjusted the p-values using FDR correction^34^.

Autocorrelation of ion images was computed using Moran’s I test available in the “moranfast” R package (https://github.com/mcooper/moranfast). The spatial correlation of ion images was computed using Pearson’s method. All correlation and autocorrelation p-values were FDR-corrected and considered significant if the p-val<0.05.

All UMAP^25^ projections were computed using the instantiation in the “uwot” R package^35^. Segmentation of UMAP projections was conducted in the “dbscan” R package ^36^ (ε=0.3). METASPACE^18^ was used for metabolite annotation against the Human Metabolome Database^22^ (10 ppm) considering all adducts available (M+, M+H, M+Na, M+K, M+NH4).

### Discovery of matrix-related signals

Using the DHB and ^13^C_6_-DHB consecutive mouse brain slices (Datasets X-Y) we compile a list of matrix-related adducts and *m/z* in three main steps: matrix-related signal discovery, contaminant-matrix adduct annotation, and analyte-matrix adduct annotation.

In the matrix-related signal discovery, all ion signals are classified into analyte, on-sample matrix-related, and off-sample matrix-related. This classification combines spatial and spectral information. The presence of all ions in each sample group (DHB on and off-sample, and ^13^C_6_-DHB on and off sample) is determined using spatial correlation and absolute mean intensity. This classification was visually validated by mapping it in a 2D UMAP projection of the spatial similarity between ions. We also compute the *m/z* shift between all possible pairs of DHB and ^13^C_6_-DHB *m/z* values (only when the ^13^C_6_ *m/z* is higher). All pairs of ions spatially classified as not analyte with an *m/z* shift matching the isotopic labeling (+6Th, +12Th, +18Th) (5 ppm) are considered matrix-related. This discovery is FDR-controlled using a bimodal decoy distribution (**μ**=N±N/2,σ=0.1). Where N corresponds to the shift used in the target search (+6Th, +12Th, +18Th).

To annotate contaminant-matrix adducts all discovered matrix-related peaks were matched against a database of theoretical DHB adducts^37–40^, generic positive-ion polarity adducts^41–44^, and DHB in-silico fragments predicted with CFM-ID^45^. This search was FDR-controlled using the decoy matrix CHCA.

Finally, to annotate analyte-matrix adducts we matched the unannotated matrix-related peaks against HMDB^22^ considering all contaminant-matrix adducts found. Exact mass searches were conducted using the R package MS2ID (https://github.com/jmbadia/MS2ID) and isotopic pattern matching was conducted using the annotation engine in rMSIcleanup^12^. This search was FDR-controlled using the decoy matrix CHCA.

### Annotation of matrix-related signals

rMSIcleanup^12^ was used to annotate matrix-related signals in Datasets C1-C14. rMSIcleanup uses the R package “enviPat”^42^ to generate the theoretical isotopic patterns of all candidate matrix-related adducts and fragments. Each candidate is matched against the experimental data and given a similarity score (S). S is the product of 3 scores: isotopic pattern similarity (S1, cosine similarity), isotopic spatial correlation (S2, weighted Pearson’s correlation), and monoisotopic ion autocorrelation (S3, Moran’s I).

The False Discovery Rate (FDR) of all annotations is estimated following a target-decoy approach^28^ using a decoy MALDI matrix. In this study, we used CHCA as a decoy matrix given its similar monoisotopic weight and structure to DHB.

rMSIcleanup uses a binary search algorithm instantiated in “Rfast”^46^ to perform efficient searches in large datasets.

## Acknowledgments

We acknowledge Dr. Denis Abu Sammour (HS Mannheim), Dr. Elisa Ruhland (IBMP), Dr. Brittney Gorman (PNNL), and Dr. Jessica Lukowski (PNNL) and respective colleagues as the original contributors of the METASPACE datasets used for validation. We acknowledge the insightful discussions with Dr. Oscar Yanes and Dr. Maria Vinaixa. Figure 3 was created with BioRender.com.

This study was funded by the Spanish Ministry of Economy and Competitivity (RTI2018096061-B-100). GB acknowledges the financial support of the European Union’s Horizon 2020 research and innovation program under the Marie Skłodowska-Curie grant agreement No. 713679 and the Universitat Rovira i Virgili (URV). MB and OB thank the financial support of the Spanish Government-MCIN, the national agency of investigation (AEI/10.13039/501100011033), and the European Regional Development Fund-ERDF (project; PID2020-120584RB-I00 to OB and FPU Fellowship; FPU19/01969 to MB). LS acknowledges the financial support of Universitat Rovira i Virgili through the pre-doctoral grant 2017PMF-PIPF-60. MGA acknowledges the financial support from the Agency for Management of University and Research Grants of the Generalitat de Catalunya (AGAUR) through the postdoctoral grant 2018 BP 00188.

## Authors’ contributions

GB, MB, MG, OB, XC, and PR developed the concept for the study. MB and OB designed, performed, and validated the synthesis of 13*C*^6^-DHB. RZ, and BH designed and performed mass spectrometry imaging experiments. GB developed the computational workflow in collaboration with LS, MG, XC, and PR. GB and MB wrote the original manuscript with substantial edits and contributions from all authors. BH, OB, and XC provided supervision, project administration, and funding.

## Data availability

Datasets A1-A4 and B1-B20 supporting the conclusions of this article are available in the Mendeley Data repository https://doi.org/10.17632/ms3365kb5p.1. Datasets C1-C15 are available at https://metaspace2020.eu/.

## Code availability

An R implementation of the computational protocol is released as a module of rMSIcleanup^12^, an open-source R package for the annotation of matrix-related signals in MSI freely available under the terms of the GNU General Public License v3.0 (https://github.com/gbaquer/rMSIcleanup).

## Abbreviations

(DHB): 2,5-Dihydroxybenzoic acid
(^13^C_6_-DHB): 2,5-Dihydroxybenzoic acid with ^13^C-labeled aromatic ring
(CHCA): alpha-Cyano-4-hydroxycinnamic acid
(9AA): 9-Aminoacridine
(DAN): 1,5-Diaminonaphthalene
(NEDC): N-(1-naphthyl) ethylenediamine dihydrochloride
(ML): Machine Learning
(AI): Artificial Intelligence
(HMDB): Human Metabolome Database
(SIL): Standard Isotope Labeled
(FDR): False Discovery Rate
(NMR): Nuclear Magnetic Resonance
(MALDI): Matrix Assisted Laser Desorption Ionization
(MSI): Mass Spectrometry Imaging
(TIC): Total Ion Current
(UMAP): Uniform Manifold Approximation and Projection
(CFM-ID): Competitive Fragmentation Modeling for Metabolite Identification
(FTICR): Fourier-transform ion cyclotron resonance
(TOF): Time Of Flight
(DBSCAN): Density-Based Spatial Clustering of Applications with Noise
(ITO): Indium tin oxide
(AUC): Area Under the Curve

## References

1. Unsihuay, D., Mesa Sanchez, D. & Laskin, J. Quantitative Mass Spectrometry Imaging of Biological Systems. Annu. Rev. Phys. Chem. 72, 307–329 (2021).

2. Chaurand, P., Norris, J. L., Cornett, D. S., Mobley, J. A. & Caprioli, R. M. New developments in profiling and imaging of proteins from tissue sections by MALDI mass spectrometry. J. Proteome Res. 5, 2889–2900 (2006).

3. Rohner, T. C., Staab, D. & Stoeckli, M. MALDI mass spectrometric imaging of biological tissue sections. Mech. Ageing Dev. 126, 177–185 (2005).

4. Alexandrov, T. Spatial Metabolomics and Imaging Mass Spectrometry in the Age of Artificial Intelligence. Annu Rev Biomed Data Sci 3, 61–87 (2020).

5. Greer, T., Sturm, R. & Li, L. Mass spectrometry imaging for drugs and metabolites. J. Proteomics 74, 2617–2631 (2011).

6. Gao, S.-Q. et al. Mass spectrometry imaging technology in metabolomics: A systematic review. Biomed. Chromatogr. e5494 (2022).

7. Notarangelo, G. et al. Oncometabolite d-2HG alters T cell metabolism to impair CD8+ T cell function. Science 377, 1519–1529 (2022).

8. Coy, S. et al. Single cell spatial analysis reveals the topology of immunomodulatory purinergic signaling in glioblastoma. Nat. Commun. 13, 4814 (2022).

9. Wang, L. et al. PARP-inhibition reprograms macrophages toward an anti-tumor phenotype. Cell Rep. 41, 111462 (2022).

10. Conage-Pough, J. E. et al. WSD-0922, a novel brain-penetrant inhibitor of EGFR, promotes survival in glioblastoma mouse models. Neuro Oncol Adv vdad066 (2023).

11. Shariatgorji, M. et al. Deuterated matrix-assisted laser desorption ionization matrix uncovers masked mass spectrometry imaging signals of small molecules. Anal. Chem. 84, 7152–7157 (2012).

12. Baquer, G. et al. rMSIcleanup: an open-source tool for matrix-related peak annotation in mass spectrometry imaging and its application to silver-assisted laser desorption/ionization. J. Cheminform. 12, 45 (2020).

13. Baquer, G. et al. What are we imaging? Software tools and experimental strategies for annotation and identification of small molecules in mass spectrometry imaging. Mass Spectrom. Rev. e21794 (2022).

14. Calvano, C. D., Monopoli, A., Cataldi, T. R. I. & Palmisano, F. MALDI matrices for low molecular weight compounds: an endless story? Anal. Bioanal. Chem. 410, 4015–4038 (2018).

15. Dong, W. et al. Phospholipid analyses by MALDI-TOF/TOF mass spectrometry using 1,5-diaminonaphthalene as matrix. Int. J. Mass Spectrom. 343-344, 15–22 (2013).

16. Ràfols, P. et al. Assessing the potential of sputtered gold nanolayers in mass spectrometry imaging for metabolomics applications. PLoS One 13, e0208908 (2018).

17. Iakab, S.-A. et al. SALDI-MS and SERS Multimodal Imaging: One Nanostructured Substrate to Rule Them Both. Anal. Chem. 94, 2785–2793 (2022).

18. Alexandrov, T. et al. METASPACE: A community-populated knowledge base of spatial metabolomes in health and disease. bioRxiv (2019) doi:10.1101/539478.

20. Janda, M. et al. Determination of Abundant Metabolite Matrix Adducts Illuminates the Dark Metabolome of MALDI-Mass Spectrometry Imaging Datasets. Anal. Chem. 93, 8399–8407 (2021).

20. Ovchinnikova, K., Kovalev, V., Stuart, L. & Alexandrov, T. OffsampleAI: artificial intelligence approach to recognize off-sample mass spectrometry images. BMC Bioinformatics 21, 129 (2020).

21. Elias, J. E. & Gygi, S. P. Target-decoy search strategy for increased confidence in large-scale protein identifications by mass spectrometry. Nat. Methods 4, 207–214 (2007).

22. Wishart, D. S. et al. HMDB 4.0: the human metabolome database for 2018. Nucleic Acids Res. 46, D608–D617 (2018).

23. Taylor, A. J., Dexter, A. & Bunch, J. Exploring Ion Suppression in Mass Spectrometry Imaging of a Heterogeneous Tissue. Anal. Chem. 90, 5637–5645 (2018).

24. Smets, T. et al. Evaluation of Distance Metrics and Spatial Autocorrelation in Uniform Manifold Approximation and Projection Applied to Mass Spectrometry Imaging Data. Anal. Chem. 91, 5706–5714 (2019).

25. McInnes, L., Healy, J. & Melville, J. UMAP: Uniform Manifold Approximation and Projection for Dimension Reduction. arXiv [stat.ML] (2018).

26. Tortorella, S. et al. LipostarMSI: Comprehensive, Vendor-Neutral Software for Visualization, Data Analysis, and Automated Molecular Identification in Mass Spectrometry Imaging. J. Am. Soc. Mass Spectrom. 31, 155–163 (2020).

27. Sementé, L., Baquer, G., García-Altares, M., Correig-Blanchar, X. & Ràfols, P. rMSIannotation: A peak annotation tool for mass spectrometry imaging based on the analysis of isotopic intensity ratios. Anal. Chim. Acta 1171, 338669 (2021).

28. Palmer, A. et al. FDR-controlled metabolite annotation for high-resolution imaging mass spectrometry. Nat. Methods 14, 57–60 (2017).

29. Shah, N. J., Worthoff, W. A. & Langen, K.-J. Imaging of sodium in the brain: a brief review. NMR Biomed. 29, 162–174 (2016).

30. Carrasco, S. & Mérida, I. Diacylglycerol, when simplicity becomes complex. Trends Biochem. Sci. 32, 27–36 (2007).

31. Schubert, E., Sander, J., Ester, M., Kriegel, H. P. & Xu, X. DBSCAN Revisited, Revisited: Why and How You Should (Still) Use DBSCAN. ACM Trans. Database Syst. 42, 1–21 (2017).

32. Ràfols, P. et al. RMSIproc: An R package for mass spectrometry imaging data processing. Bioinformatics 36, 3618–3619 (2020).

33. Pinheiro, J. nlme : Linear and nonlinear mixed effects models. R package version 3.1–96. http://cran.r-project.org/web/packages/nlme/ (2009).

34. Benjamini, Y. & Hochberg, Y. Controlling the false discovery rate: A practical and powerful approach to multiple testing. J. R. Stat. Soc. 57, 289–300 (1995).

35. Melville, J., Lun, A. & Djekidel, M. uwot: the Uniform Manifold Approximation and Projection (UMAP) method for dimensionality reduction. R package version 0.1. 8. Preprint at (2020).

36. Hahsler, M., Piekenbrock, M. & Doran, D. dbscan: Fast density-based clustering with R. J. Stat. Softw. 91, 1–30 (2019).

37. Keller, B. O. & Li, L. Discerning matrix-cluster peaks in matrix-assisted laser desorption/ionization time-of-flight mass spectra of dilute peptide mixtures. J. Am. Soc. Mass Spectrom. 11, 88–93 (2000).

38. Bourcier, S., Bouchonnet, S. & Hoppilliard, Y. Ionization of 2,5-dihydroxybenzoic acid (DHB) matrix-assisted laser desorption ionization experiments and theoretical study. International Journal of Mass Spectrometry vols 210-211 59–69 Preprint at https://doi.org/10.1016/s1387-3806(01)00446-8 (2001).

39. Wallace, W. E., Arnould, M. A. & Knochenmuss, R. 2,5-Dihydroxybenzoic acid: laser desorption/ionisation as a function of elevated temperature. International Journal of Mass Spectrometry vol. 242 13–22 Preprint at https://doi.org/10.1016/j.ijms.2004.11.011 (2005).

40. Petković, M. et al. Detection of Adducts with Matrix Clusters in the Positive and Negative Ion Mode MALDI-TOF Mass Spectra of Phospholipids. Zeitschrift für Naturforschung B 64, 331–334 (2009).

41. Strohalm, M., Kavan, D., Novák, P., Volný, M. & Havlícek, V. mMass 3: a cross-platform software environment for precise analysis of mass spectrometric data. Anal. Chem. 82, 4648–4651 (2010).

42. Loos, M., Gerber, C., Corona, F., Hollender, J. & Singer, H. Accelerated isotope fine structure calculation using pruned transition trees. Anal. Chem. 87, 5738–5744 (2015).

43. Huang, N., Siegel, M. M., Kruppa, G. H. & Laukien, F. H. Automation of a Fourier transform ion cyclotron resonance mass spectrometer for acquisition, analysis, and e-mailing of high-resolution exact-mass electrospray ionization mass spectral data. Journal of the American Society for Mass Spectrometry vol. 10 1166–1173 Preprint at https://doi.org/10.1016/s1044-0305(99)00089-6 (1999).

44. Keller, B. O., Sui, J., Young, A. B. & Whittal, R. M. Interferences and contaminants encountered in modern mass spectrometry. Anal. Chim. Acta 627, 71–81 (2008).

45. Wang, F. et al. CFM-ID 4.0: More Accurate ESI-MS/MS Spectral Prediction and Compound Identification. Anal. Chem. 93, 11692–11700 (2021).

46. Papadakis, M., Tsagris, M., Dimitriadis, M. & Fafalios, S. Rfast: A collection of efficient and extremely fast R functions. R package version.

