## Supplementary Materials for "Discovering Matrix Adducts for Enhanced Metabolite Profiling with Stable Isotope-Labeled MALDI-MSI"

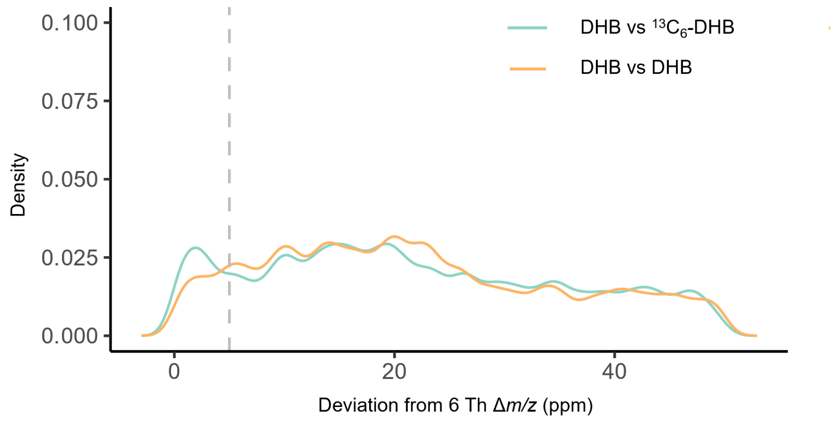

**Supplementary Figure 1.** Density distribution of ppm error of +2 DA *m/z* shift.

**Supplementary Table 1.** List of the 24 MALDI MSI datasets used for method development and validation. Sample type, sample preparation, and MALDI-MSI acquisition parameters.

| No. | Species | Tissue type | Matrix deposition | Lateral Res. (um) | *m/z* range | Ion source / Mass spectrometer | Acq. Mode | Notes | Ref. / Data availability |
|---|---|---|---|---|---|---|---|---|---|
| A1-A2 | *Mus musculus* | Brain | DHB, TM sprayer | 100 µm | 100-1000 | 9.4T Solarix FTICR (Bruker) | Positive/Profile | Technical replicates | This study https://doi.org/10.17632/ms3365kb5p.1 |
| A3-A4 | *Mus musculus* | Brain | $13C^6$-DHB, TM sprayer | 100 µm | 100-1000 | 9.4T Solarix FTICR (Bruker) | Positive/Profile | Technical replicates | This study https://doi.org/10.17632/ms3365kb5p.1 |
| B1-B2 | *Mus musculus* | Brain | 9AA, TM sprayer | 100 µm | 100-1000 | 9.4T Solarix FTICR (Bruker) | Negative/Profile | Technical replicates | This study https://doi.org/10.17632/ms3365kb5p.1 |
| B3-B4 | - | Matrix control | 9AA, TM sprayer | 100 µm | 100-1000 | 9.4T Solarix FTICR (Bruker) | Negative/Profile | Technical replicates | This study https://doi.org/10.17632/ms3365kb5p.1 |
| B5-B6 | *Mus musculus* | Brain | NEDC, TM sprayer | 100 µm | 100-1000 | 9.4T Solarix FTICR (Bruker) | Negative/Profile | Technical replicates | This study https://doi.org/10.17632/ms3365kb5p.1 |
| B7-B8 | - | Matrix control | NEDC, TM sprayer | 100 µm | 100-1000 | 9.4T Solarix FTICR (Bruker) | Negative/Profile | Technical replicates | This study |

| | | | | | | | | | |
|---|---|---|---|---|---|---|---|---|---|
| | | | | | | | | | https://doi.org/10.17632/ms3365kb5p.1 |
| B9-B10 | *Mus musculus* | Brain | NOR, TM sprayer | 100 µm | 100-1000 | 9.4T Solarix FTICR (Bruker) | Negative/Profile | Technical replicates | This study<br><br>https://doi.org/10.17632/ms3365kb5p.1 |
| B11-B12 | - | Matrix control | NOR, TM sprayer | 100 µm | 100-1000 | 9.4T Solarix FTICR (Bruker) | Negative/Profile | Technical replicates | This study<br><br>https://doi.org/10.17632/ms3365kb5p.1 |
| B13-B14 | *Mus musculus* | Brain | Au, TM sprayer | 100 µm | 100-1000 | 9.4T Solarix FTICR (Bruker) | Positive/Profile | Technical replicates | This study<br><br>https://doi.org/10.17632/ms3365kb5p.1 |
| B15-B16 | - | Matrix control | Au, TM sprayer | 100 µm | 100-1000 | 9.4T Solarix FTICR (Bruker) | Positive/Profile | Technical replicates | This study<br><br>https://doi.org/10.17632/ms3365kb5p.1 |
| B17-B18 | *Mus musculus* | Brain | DHB, TM sprayer | 100 µm | 100-1000 | 9.4T Solarix FTICR (Bruker) | Positive/Profile | Technical replicates | This study<br><br>https://doi.org/10.17632/ms3365kb5p.1 |
| B19-B20 | - | Matrix control | DHB, TM sprayer | 100 µm | 100-1000 | 9.4T Solarix FTICR (Bruker) | Positive/Profile | Technical replicates | This study<br><br>https://doi.org/10.17632/ms3365kb5p.1 |

**Supplementary Table 2.** List of the 14 METASPACE (Alexandrov et al. 2019) MALDI MSI datasets used for validation. Sample type, sample preparation, and MALDI-MSI acquisition parameters.

| **No.** | **Species** | **Tissue type** | **Matrix deposition** | **Lateral Res. (um)** | ***m/z* range** | **Mass spectrometer** | **Acq. Mode** | **Contributor** | **Ref.** |
|---|---|---|---|---|---|---|---|---|---|
| C1-C2 | *Homo Sapiens* | Liver | DHB, TM sprayer | N/A | 150-1850 | FTICR | Positive/Centroid | Denis Abu Sammour (HS Mannheim) | (Alexandrov et al. 2019) |
| C3-C4 | *Homo Sapiens* | Brain | DHB, TM sprayer | N/A | 100-1150 | FTICR | Positive/Centroid | Elisa Ruhland (IBMP) | (Alexandrov et al. 2019) |
| C5-C7 | *Homo Sapiens* | Lung | DHB, HTX M5 Sprayer | 35 µm | 200-1200 | FTICR | Positive/Centroid | Brittney Gorman (PNNL) | (Alexandrov et al. 2019) |
| C8-C10 | *Homo Sapiens* | Kidney | DHB, TM sprayer | 35 µm | 200-1300 | FTICR | Positive/Centroid | Jessica Lukowski (PNNL) | (Alexandrov et al. 2019) |

**Supplementary Table 3.** MALDI-matrix composition for datasets A1-A4 and B1-B20.

| **Matrices** | **DHB** | **13C-DHB** | **9-AA** | **NEDC** | **NOR** |
|---|---|---|---|---|---|
| Concentration (mg/mL) | 20 | 20 | 10 | 7 | 7 |
| ACN (%) | 50 | 50 | | | |
| MeOH (%) | | | 70 | 70 | 33 |
| MilliQ (%) | 49.9 | 49.9 | 30 | 30 | |
| TFA (%) | 0.1 | 0.1 | | | |
| Chloroform (%) | | | | | 67 |

**Supplementary Table 4.** MALDI-matrix SunCollect deposition.

| SunCollect | DHB | 13C-DHB | 9-AA | NEDC | NOR |
|---|---|---|---|---|---|
| Line distance (mm) | 2 | 2 | 1 | 1 | 2 |
| Z-height (mm) | 25 | 25 | 25 | 35 | 20 |
| Layers (#) | 8 | 8 | 10 | 6 | 15 |
| #1 (µL/min) | 13 | 13 | 13 | 13 | 13 |
| #2 (µL/min) | 20 | 20 | 13 | 13 | 20 |
| #3 (µL/min) | 30 | 30 | 13 | 30 | 30 |
| #4+ (µL/min) | 40 | 40 | 13 | 40 | 40 |
| Speed XY (mm/min) | 900 | 900 | 900 | 900 | 900 |
| N2 (psi) | 35 | 35 | 35 | 35 | 35 |

**Supplementary Table 5.** Measurement polarity, laser intensity, number of shots per pixel, and laser frequency.

| **Dataset** | **A1-A2** | **A3-A4** | **B1-B4** | **B5-B8** | **B9-B12** | **B13-B16** | **B17-B20** |
|---|---|---|---|---|---|---|---|
| matrix | DHB | 13C-DHB | 9-AA | NEDC | NOR | AU | DHB |
| magnet | 12 T | 12 T | 9.4 T | 9.4 T | 9.4 T | 9.4 T | 9.4 T |
| polarity | POS | POS | NEG | NEG | NEG | POS | POS |
| m/z low | 98.3 | 98.3 | 101.1 | 101.1 | 101.1 | 101.1 | 101.1 |
| m/z high | 1000 | 1000 | 1000 | 1000 | 1000 | 1000 | 1000 |
| TD | 512k | 512k | 512k | 512k | 512k | 512k | 512k |
| laser | 35 | 35 | 65 | 80 | 75 | 65 | 60 |
| shots | 100 | 100 | 700 | 500 | 200 | 100 | 500 |
| freq | 500 | 500 | 500 | 500 | 2000 | 2000 | 2000 |
| focus | small | small | small | small | small | minimum | small |
| raster | 100x100 | 100x100 | 100x100 | 100x100 | 100x100 | 100x100 | 100x100 |

**Supplementary Table 6.** Complete list of discovered endogenous matrix adducts.

| Experimental *m/z* | Adduct | FDR | Confidence level |
|---|---|---|---|
| 177.0159 | DHB+M | 0.138462 | + |
| 183.4817 | DHB+M | 0.138462 | + |
| 184.4795 | DHB+M | 0.138462 | + |
| 188.493 | DHB+M | 0.138462 | + |
| 192.9901 | DHB+M | 0.138462 | + |
| 196.1071 | DHB+M | 0.138462 | + |
| 196.4796 | DHB+M | 0.138462 | + |
| 196.9809 | DHB+M | 0.138462 | + |
| 197.4783 | DHB+M | 0.138462 | + |
| 212.9568 | DHB+M | 0.138462 | + |
| 229.9384 | DHB+M | 0.138462 | + |
| 232.9446 | DHB+M | 0.138462 | + |
| 240.1235 | DHB+M | 0.138462 | + |
| 241.1265 | DHB+M | 0.138462 | + |
| 268.9312 | DHB+M | 0.138462 | + |

| | | | |
|---|---|---|---|
| 278.0569 | DHB+M | 0.138462 | + |
| 295.0232 | DHB+M | 0.138462 | + |
| 302.0808 | DHB+M | 0.138462 | + |
| 310.9976 | DHB+M | 0.138462 | + |
| 313.0342 | DHB+M | 0.138462 | + |
| 316.0013 | DHB+M | 0.138462 | + |
| 317.017 | DHB+M | 0.138462 | + |
| 322.0959 | DHB+M | 0.138462 | + |
| 328.9657 | DHB+M | 0.138462 | + |
| 332.8965 | DHB+M | 0.138462 | + |
| 359.0081 | DHB+M | 0.138462 | + |
| 360.0704 | DHB+M | 0.138462 | + |
| 362.0856 | DHB+M | 0.138462 | + |
| 366.9659 | DHB+M | 0.138462 | + |
| 382.8307 | DHB+M | 0.138462 | + |
| 390.1007 | DHB+M | 0.138462 | + |
| 391.967 | DHB+M | 0.138462 | + |
| 392.9629 | DHB+M | 0.138462 | + |

| | | | |
|---|---|---|---|
| 393.9193 | DHB+M | 0.138462 | + |
| 409.0614 | DHB+M | 0.138462 | + |
| 414.0162 | DHB+M | 0.138462 | + |
| 418.3054 | DHB+M | 0.138462 | + |
| 420.9163 | DHB+M | 0.138462 | + |
| 430.956 | DHB+M | 0.138462 | + |
| 470.036 | DHB+M | 0.138462 | + |
| 474.0575 | DHB+M | 0.138462 | + |
| 490.2457 | DHB+M | 0.138462 | + |
| 497.0368 | DHB+M | 0.138462 | + |
| 544.0335 | DHB+M | 0.138462 | + |
| 585.0817 | DHB+M | 0.138462 | + |
| 685.3169 | DHB+M | 0.138462 | + |
| 738.2404 | DHB+M | 0.138462 | + |
| 763.0849 | DHB+M | 0.138462 | + |
| 831.9886 | DHB+M | 0.138462 | + |
| 869.0752 | DHB+M | 0.138462 | + |
| 883.144 | DHB+M | 0.138462 | + |

| | | | |
|---|---|---|---|
| 907.487 | DHB+M | 0.138462 | + |
| 923.1411 | DHB+M | 0.138462 | + |
| 153.5236 | DHB+M | 0.138462 | + |
| 196.4796 | DHB+M | 0.138462 | + |
| 311.5289 | DHB+M | 0.138462 | + |
| 394.7467 | DHB+M | 0.138462 | + |
| 406.7492 | DHB+M | 0.138462 | + |
| 427.3117 | DHB+M | 0.138462 | + |
| 648.0488 | DHB+M | 0.138462 | + |
| 923.1411 | DHB+M | 0.138462 | + |
| 941.1575 | DHB+M | 0.138462 | + |
| 340.9644 | 2DHB+M | 0.121951 | + |
| 348.9723 | 2DHB+M | 0.121951 | + |
| 360.0181 | 2DHB+M | 0.121951 | + |
| 361.0143 | 2DHB+M | 0.121951 | + |
| 364.9479 | 2DHB+M | 0.121951 | + |
| 375.0089 | 2DHB+M | 0.121951 | + |
| 384.9754 | 2DHB+M | 0.121951 | + |

| | | | |
|---|---|---|---|
| 391.967 | 2DHB+M | 0.121951 | + |
| 392.9629 | 2DHB+M | 0.121951 | + |
| 394.9211 | 2DHB+M | 0.121951 | + |
| 394.9617 | 2DHB+M | 0.121951 | + |
| 406.9603 | 2DHB+M | 0.121951 | + |
| 430.956 | 2DHB+M | 0.121951 | + |
| 481.2172 | 2DHB+M | 0.121951 | + |
| 488.0139 | 2DHB+M | 0.121951 | + |
| 489.0531 | 2DHB+M | 0.121951 | + |
| 528.9874 | 2DHB+M | 0.121951 | + |
| 540.9662 | 2DHB+M | 0.121951 | + |
| 544.0335 | 2DHB+M | 0.121951 | + |
| 560.0065 | 2DHB+M | 0.121951 | + |
| 571.9807 | 2DHB+M | 0.121951 | + |
| 603.091 | 2DHB+M | 0.121951 | + |
| 668.0517 | 2DHB+M | 0.121951 | + |
| 685.0229 | 2DHB+M | 0.121951 | + |
| 685.3169 | 2DHB+M | 0.121951 | + |

| | | | |
|---|---|---|---|
| 808.8479 | 2DHB+M | 0.121951 | + |
| 810.0001 | 2DHB+M | 0.121951 | + |
| 977.0996 | 2DHB+M | 0.121951 | + |
| 979.1153 | 2DHB+M | 0.121951 | + |
| 986.1133 | 2DHB+M | 0.121951 | + |
| 323.5466 | 2DHB+M | 0.121951 | + |
| 628.0607 | 2DHB+M | 0.121951 | + |
| 694.9814 | 2DHB+M | 0.121951 | + |
| 816.0104 | 2DHB+M | 0.121951 | + |
| 978.0475 | 2DHB+M | 0.121951 | + |
| 471.0412 | 3DHB+M | 0.28125 | + |
| 473.972 | 3DHB+M | 0.28125 | + |
| 497.0368 | 3DHB+M | 0.28125 | + |
| 511.032 | 3DHB+M | 0.28125 | + |
| 530.985 | 3DHB+M | 0.28125 | + |
| 537.032 | 3DHB+M | 0.28125 | + |
| 570.9783 | 3DHB+M | 0.28125 | + |
| 581.2709 | 3DHB+M | 0.28125 | + |

| | | | |
|---|---|---|---|
| 584.9589 | 3DHB+M | 0.28125 | + |
| 614.9233 | 3DHB+M | 0.28125 | + |
| 842.0349 | 3DHB+M | 0.28125 | + |
| 275.7525 | 3DHB+M | 0.28125 | + |
| 329.5111 | 3DHB+M | 0.28125 | + |
| 529.9907 | 3DHB+M | 0.28125 | + |
| 555.9419 | 3DHB+M | 0.28125 | + |
| 587.5494 | 3DHB+M | 0.28125 | + |
| 612.079 | 3DHB+M | 0.28125 | + |
| 716.0391 | 3DHB+M | 0.28125 | + |
| 879.9781 | 3DHB+M | 0.28125 | + |
| 978.0475 | 3DHB+M | 0.28125 | + |

**Supplementary Table 7.** List of discovered endogenous matrix adducts matched against HMDB.

| Theoretical m/z | Experimental m/z | Error (ppm) | Endogenous compound | DHB adduct | FDR | Confidence level |
|---|---|---|---|---|---|---|
| 177.0164 | 177.0159 | 2.92 | Na | DHB | 13.85% | +++ |
| 177.0164 | 177.0159 | 2.92 | H2O | DHB+Na-H2O | 13.85% | +++ |
| 192.9903 | 192.9901 | 1.05 | K | DHB | 13.85% | +++ |

| | | | | | | |
|---|---|---|---|---|---|---|
| 192.9903 | 192.9901 | 1.05 | H2O | DHB+K-H2O | 13.85% | +++ |
| 196.1079 | 196.1071 | 4.13 | C5H11NO2 | DHB+H2 | 13.85% | +++ |
| 232.9449 | 232.9446 | 1.56 | Br | DHB | 13.85% | +++ |
| 240.1236 | 240.1235 | 0.36 | C3H8 | DHB+C2H3N+H | 13.85% | +++ |
| 240.1236 | 240.1235 | 0.36 | C5H12N | DHB | 13.85% | +++ |
| 240.1236 | 240.1235 | 0.36 | C6H14NO2 | DHB-H2O-CO | 13.85% | +++ |
| 295.0220 | 295.0232 | -3.88 | C4H8O4 | DHB+K-H2O | 13.85% | +++ |
| 322.0959 | 322.0959 | -0.06 | C10H17NOS | DHB+2Na | 13.85% | +++ |
| 328.9664 | 328.9657 | 2.21 | Cs | DHB+C2H3N+H | 13.85% | +++ |
| 332.8957 | 332.8965 | -2.21 | Ce | DHB+K | 13.85% | +++ |
| 359.0073 | 359.0081 | -2.26 | C6H6O6S | DHB-H | 13.85% | +++ |
| 359.0072 | 359.0081 | -2.49 | C6H13O7PS | DHB+H+Na | 13.85% | +++ |
| 360.0719 | 360.0704 | 4.18 | C10H7NO4 | DHB+H | 13.85% | +++ |
| 360.0695 | 360.0704 | -2.50 | C6H6O4 | DHB+C2H3N+Na | 13.85% | +++ |
| 360.0720 | 360.0704 | 4.43 | C8H6ClNO3 | DHB+C3H8O+Na+H | 13.85% | +++ |
| 360.0693 | 360.0704 | -3.27 | C7H10O7 | DHB | 13.85% | +++ |
| 360.0719 | 360.0704 | 4.18 | C6H4O4 | DHB+2C2H3N+H | 13.85% | +++ |
| 362.0852 | 362.0856 | -1.24 | C6H8O4 | DHB+C2H3N+Na | 13.85% | +++ |

| | | | | | | |
|---|---|---|---|---|---|---|
| 362.0849 | 362.0856 | -2.00 | C7H14O7 | DHB-2H | 13.85% | +++ |
| 362.0851 | 362.0856 | -1.57 | C13H15NO2 | DHB-H+3Na | 13.85% | +++ |
| 392.9647 | 392.9629 | 4.51 | C5FeN6O | DHB+Na | 13.85% | +++ |
| 392.9621 | 392.9629 | -2.14 | ClTl | DHB-H | 13.85% | +++ |
| 409.0621 | 409.0614 | 1.67 | C6H9NO4 | DHB+C2HF3O2-H2O | 13.85% | +++ |
| 409.0617 | 409.0614 | 0.81 | C12H14O8 | DHB+2Na | 13.85% | +++ |
| 470.0344 | 470.0360 | -3.22 | C6H11O6PS2 | DHB+C2H3N+H | 13.85% | +++ |
| 474.0568 | 474.0575 | -1.66 | C9H18NO8P | DHB+K-H2O | 13.85% | +++ |
| 490.2441 | 490.2457 | -3.28 | C17H26O4 | DHB+C2H3N+H | 13.85% | +++ |
| 497.0364 | 497.0368 | -0.96 | C10H13N2O7P | DHB+K | 13.85% | +++ |
| 497.0390 | 497.0368 | 4.29 | C12H16O12 | DHB-H+3Na | 13.85% | +++ |
| 585.0790 | 585.0817 | -4.56 | C19H18N6O5S3 | DHB+H2 | 13.85% | +++ |
| 685.3141 | 685.3169 | -4.17 | C30H52O7P2 | DHB+H+Na | 13.85% | +++ |
| 907.4885 | 907.4870 | 1.70 | C39H72N2O15 | DHB+H+Na | 13.85% | +++ |
| 648.0473 | 648.0488 | -2.37 | C16H17N5O7S2 | DHB+K | 13.85% | +++ |
| 941.1581 | 941.1575 | 0.68 | C22H31N9O13P3S | DHB+CH3OH+H | 13.85% | +++ |
| 267.1318 | 267.1324 | -2.31 | C3H7N | 2DHB+3H2O+2H | 12.20% | +++ |
| 430.9559 | 430.9560 | -0.23 | Ce | 2DHB+H-H2O | 12.20% | +++ |

| | | | | | | |
|---|---|---|---|---|---|---|
| 488.0155 | 488.0139 | 3.32 | C11H14O7S | 2DHB+K+Na-H2O | 12.20% | +++ |
| 489.0548 | 489.0531 | 3.32 | C5H8N2O3 | 2DHB-2H+K | 12.20% | +++ |
| 489.0548 | 489.0531 | 3.32 | C5H8N2O4 | 2DHB+K-H2O | 12.20% | +++ |
| 544.0334 | 544.0335 | -0.10 | C5H11NO3Se | 2DHB+Na | 12.20% | +++ |
| 544.0314 | 544.0335 | -3.70 | C10H7Cl2N3O2 | 2DHB-2H2O+H | 12.20% | +++ |
| 560.0073 | 560.0065 | 1.52 | C5H11NO3Se | 2DHB+K | 12.20% | +++ |
| 603.0905 | 603.0910 | -0.80 | C15H12O4 | 2DHB+K | 12.20% | +++ |
| 603.0897 | 603.0910 | -2.13 | C8H9NO5S | 2DHB+C2H3N+Na | 12.20% | +++ |
| 603.0905 | 603.0910 | -0.80 | C15H14O4 | 2DHB-2H+K | 12.20% | +++ |
| 603.0905 | 603.0910 | -0.80 | C15H14O5 | 2DHB+K-H2O | 12.20% | +++ |
| 685.3200 | 685.3169 | 4.47 | C21H40O5 | 2DHB+Na-H2O | 12.20% | +++ |
| 809.9966 | 810.0001 | -4.37 | C15H14INO7S | 2DHB+Na | 12.20% | +++ |
| 977.1026 | 977.0996 | 3.10 | C28H24O17 | 2DHB-2H+K | 12.20% | +++ |
| 979.1183 | 979.1153 | 3.03 | C28H24O17 | 2DHB+K | 12.20% | +++ |
| 628.0584 | 628.0607 | -3.54 | C6H8O9S | 2DHB+C2H3N+Na | 12.20% | +++ |
| 816.0123 | 816.0104 | 2.39 | C10H15ClN5O12P3 | 2DHB+H-H2O | 12.20% | +++ |
| 816.0123 | 816.0104 | 2.32 | C11H16N5O11P3 | 2DHB+K-H2O | 12.20% | +++ |
| 419.0403 | 419.0420 | -3.85 | C10H8O | 3DHB+K+Na-H2O | 28.13% | +++ |

| | | | | | | |
|---|---|---|---|---|---|---|
| 529.9889 | 529.9907 | -3.49 | Rb | 3DHB+H-H2O | 28.13% | +++ |
| 529.9886 | 529.9907 | -3.95 | Cu | 3DHB+Na-H2O | 28.13% | +++ |
